# Transcriptomes Often Resemble Biological Aging Upon Perturbation

**DOI:** 10.1101/2023.03.28.534472

**Authors:** Thomas Stoeger

## Abstract

Aging is deeply interwoven with cellular and physiological biology, but the extent is not known. Here, I demonstrate that the Gene Length-dependent Transcription Decline (GLTD) which marks age-dependent transcriptional change is not specific to aging. Instead, it follows perturbations in 5,231 of 11,959 public gene expression profiles. My analysis suggests the hypothesis that during the passage of time, exposure to myriad perturbations change transcriptomes in a way that would be described as “aging”.

## Main Text

Conditions that promote health tend to delay phenotypic manifestations of aging, whereas the inverse tends to hold for conditions that worsen health^1^. However, it is not known to what extent perturbations of cellular and organismal physiology yield phenotypes resembling those encountered during aging^2^.

To address this question, I drew from the recent insight that transcriptomic changes in humans and mice with age can be approximated by a negative correlation between gene length and transcriptomic change^3-7^, termed Gene Length-dependent Transcription Decline (GLTD) [*in press, Trends in Genetics*^*8*^]. Unlike most molecular and physiological markers of age^9^, GLTD can be retrospectively measured in a wide set of biomedical studies, as transcriptomics are frequently reported in a wide vary of biomedical settings. I begin my investigation with resources that aggregate studies.

EBI Gene Expression Atlas (EBI-GXA) gathers and processes high-quality transcriptomic studies^10^, which reflect areas of interest to individual investigators (10 of 10 randomly sampled studies, Extended Data Table 1). I considered those 7,108 (4,077 human and 3,031 mouse) differential gene expression profiles that report the relative fold-changes of transcripts of perturbed samples relative to control samples for at least 10,000 genes. Because GLTD in aging originates in a loss of productive transcription over the gene body^7^, I quantified gene length through the median length of primary transcripts, omitting non-coding regulatory regions. As before^5^, I quantified length-correlated change through the Spearman correlation between gene length and relative fold-changes of transcripts of protein-coding genes using Kendall and Stuart’s method for *P* values^11^.

Though EBI-GXA does not directly report age, it reproduces insights into a model of accelerated aging^12^ in which GLTD has been firmly established^3,7^. Specifically, mice with mutations in the DNA excision repair protein *DNA excision repair protein ERCC-1* (ERCC1) gene show a significant negative correlation (Figure 1A, Spearman correlation: -0.14, *P* value = 2.4E-68). Further, the 7^th^ most negative correlation in EBI-GXA derives from UV-irradiated mice with mutations in the related *DNA repair protein complementing XP-A cells* (XPA) gene and ERCC6 genes (Figure 1B; Spearman correlation: -0.35, *P* value of “∼ 0” as not precisely computable). Complete results are provided as Extended Data Table 2.

**Figure 1.**
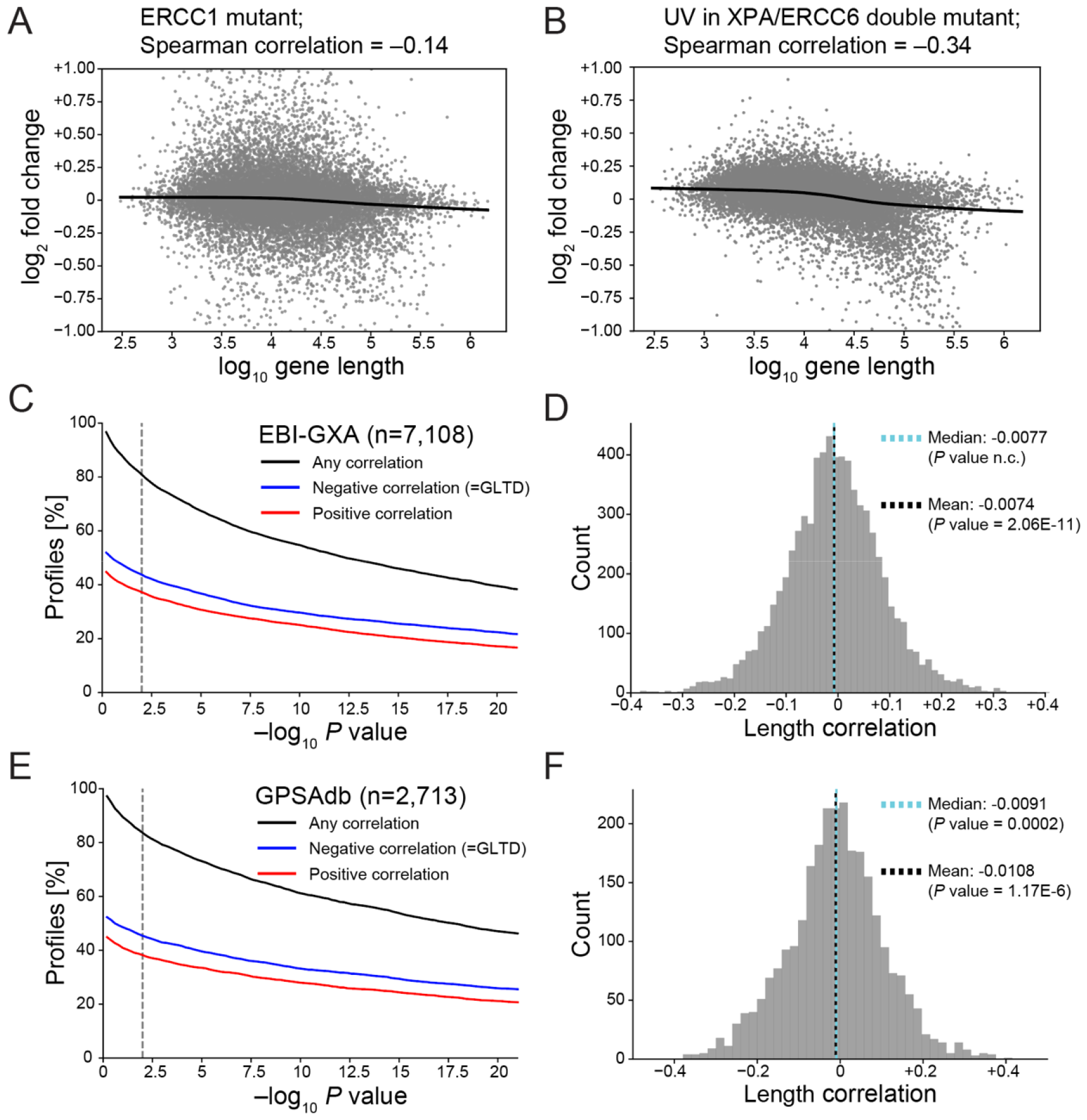
Gene Length-dependent Transcription Decline (GLTD) is frequent in public transcriptomic data. (A, B) Transcriptomic fold changes of genes (grey dots). (C) Fraction of gene expression profiles with length-correlated change in EBI-GXA^10^ at different *P* value thresholds, determined by Kendall and Stuart’s method^11^. Dashed vertical line indicates *P* value of 0.01. (D) Histogram of all observed length-correlated changes in EBI-GXA, including non-significant ones. *P* value of median inferred from bootstrapping and *P* value of mean from two-sided one group T-test. n.c. is non computable as close to 0. (E, F) as (A,B) but for GPSAdb^19^.

In 44% of all profiles, I found that perturbations often resemble biological aging, as they often yield a negative correlation (=GLTD) (Figure 1C, at a significance cutoff of *P* value = 0.01). These perturbations were not confined to any single domain of biology (Figure 1C, Extended Data Table 2). Importantly, perturbations preferentially caused length-correlated changes in the direction that is dominant in aging^3-7^, as more profiles showed a negative correlation (=GLTD) than positive correlation (Figure 1C, Extended Data Figure 2, e.g.: Fisher’s Exact *P* value of 1.76E-5 when for length correlations considering a *P* value threshold of 0.01).

Similarly, the median and mean effect size of length-correlated change both were negative (P∼”0” via bootstrapping, and P=2.06E-11 via T-test, respectively). While effect sizes were small (−0.0077 and -0.0074, respectively), the negative effect sizes include all perturbations in EBI-GXA independent of the significance of length-correlated change, consistent with the finding that age-dependent molecular change often^13,14^ (but not always^14,15^) appears to be a gradual process in which individual steps are not readily discernable from each other.

As in aging^5^, GLTD following perturbations was largely independent of transcriptional regulation of known age-associated pathways, as 40% of profiles still displayed GLTD when excluding 7,691 of 19,163 human and 8,212 of 21,167 mouse protein-coding genes that have been annotated by the Gene Ontology Consortium^16^ to participate in any of 27 processes regulated by age^17^, some as broad as “metabolism”, “inflammation”, “synaptic transmission”, and “stress” (Extended Data Figure 2).

While length-correlated changes in gene expression profiles have been suspected to have technical origins in RNA-sequencing (RNA-seq)^18^, negative correlations (=GLTD) are also more numerous than positive correlations in human and mouse microarray data (Extended Data Figure 3). While there is no discernable preference for positive or negative correlations in RNA-seq data of humans or mice (Extended Data Figure 4), this result may be attributable to lower statistical power (n=486 and n=371 versus n=2,788 and n=1,988 profiles) or increased technical variability. Length-correlated changes in RNA-seq studies have been suspected to possibly originate from length-dependent technical artifacts^18^. However, the preferential directionality of the correlations favors biological origins, and this manuscript will subsequently confirm this directionally through a variety of experimental methods and datasets, and observations on nascent transcripts.

Another resource, GPSAdb^19^, that exclusively contains RNA-seq data following loss-of-function perturbations of individual genes, confirms that GLTD is common after perturbations. GPSAdb aggregates a total of 2,713 profiles mapping to unambiguous gene identifiers, corresponding to 1,249 genes. Among these profiles, 45% show GLTD at a *P* value of 0.01 (Figure 1E, Extended Data Table 3). Again, there are more negative than positive correlations (Figure 1E and Extended Data Figure 5, e.g.: Fisher’s Exact *P* value of 0.004 when for length correlations considering a *P* value threshold of 0.01), and the median and mean effect size of length-correlated change are negative (Figure 1F, bootstrapped *P* value 0.0002; Figure 1F, *P* value of 1.17E-6, T-test).

Closer inspection of GPSAdb hints toward further complexity. Loss-of-function by RNA interferences (n=1,844) or genetic knockout (n=487) yielded median length correlations that are negative, but CRISPR interferences (CRISPRi) (n=73) yielded median length correlations that are positive (+0.09, bootstrapped P value of ∼0 as not precisely computable). While the lower number of perturbations with CRISPRi may have increased the chance for uncontrolled covariates, the reasons the observed increase are presently unexplained.

Avoiding potential confounding factors of data aggregation and more comprehensively inspecting the loss-of-function of genes, I next reanalyze one recent perturb-seq study^20^ that combined perturbations in a single experiment rather than after parallel experiments. The authors applied CRISPRi constructs against all expressed genes in K562 cells, identified 1,958 genes to cause a strong transcriptional phenotype, and provide bioinformatically processed data for those perturbations.

42% of profiles showed GLTD at a *P* value of 0.01 (Figure 2A, Extended Data Table 4). Opposing the observations of CRISPRi in GPSAdb, more perturbations yielded a negative correlation (=GLTD) rather than positive correlation (Figure 2B and Extended Data Figure 7, e.g.: Fisher’s Exact *P* value of 6.3E-38 when for length correlations considering a P value threshold of 0.01). The median and mean effect sizes of length-correlated change again were negative (Figure 2B, bootstrapped *P* value of ∼0 as not precisely computable; and P value of 8.31E-79, T-test, respectively).

**Figure 2.**
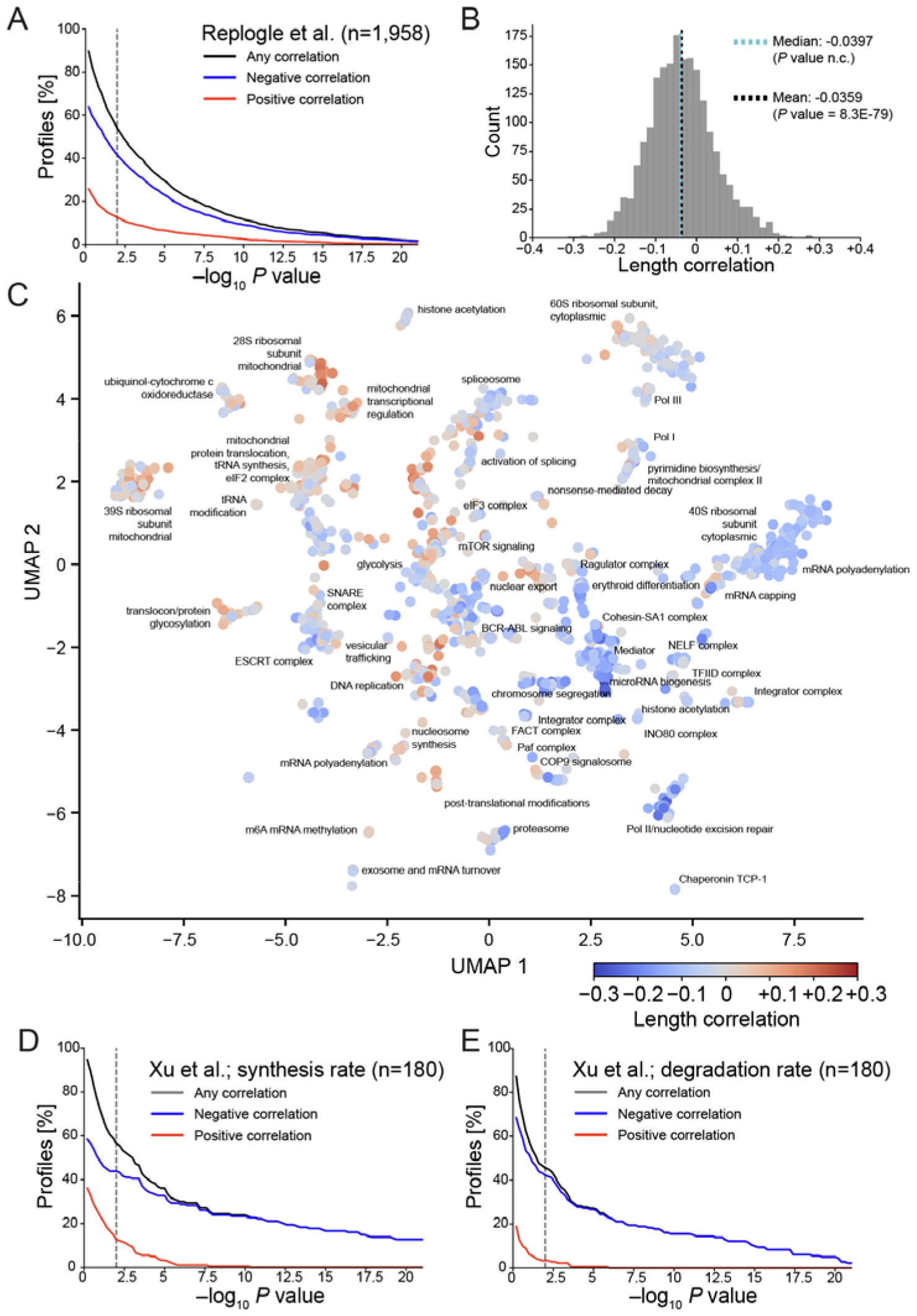
Gene Length-dependent Transcription Decline (GLTD) is frequent following the loss of genes. (A) Fraction of gene expression profiles with length-correlated change in Replogle et al.^20^ at different *P* value thresholds, determined by Kendall and Stuart’s method^11^. Dashed vertical line indicates *P* value of 0.01. (B) Histogram of all observed length-correlated changes in in Replogle et al., including non-significant ones. *P* value of median inferred from bootstrapping and *P* value of mean from two-sided one group T-test. n.c. is non computable as close to 0. (C) Length-correlated change superimposed on map by Replogle et al., a Uniform Manifold Approximation and Projection (UMAP), of genes according to the effect of their loss-of-function toward the transcriptome. (D) As (B) but for synthesis rates contained in Xu et al.^21^. (E) As (B) but for synthesis rates contained in Xu et al.^21^.

Superimposing length-correlated change onto a map created by the authors of this datasets to depict global dependencies of cellular transcriptomes on the function of individual genes identified length-correlated change as a latent factor aligning with the first two dimensions of the map (Spearman correlations of -0.40 and +0.30; P values of 1.2E-77 and 2.0E-42, respectively). This alignment does not reflect the length of the genes whose function is lost (Extended Data Figure 8A), is not affected by excluding differentially expressed genes annotated the for any of 27 processes associated with aging^17^ (Extended Data Figure 8B), and holds when independently mapping genes through Principal Components (Extended Data Figure 8C). Resembling observations in aging in which known transcription factors are less informative than gene length^5^, length-correlated change better aligns with the map than the maximal alignment observed for any of 1,268 transcription factors (Extended Data Table 5 with maximal effect sizes of correlation of +0.11 and +0.13, respectively). Gene Ontology Enrichment analysis confirmed the preceding annotation that accompanied the map^20^ (Figure 2C), as GLTD is most enriched among genes that contribute to transcription or translation (Extended Data Table 6).

To provide initial mechanistic clues, I turn toward a recent parallel perturbation experiment^21^ that succeeded in biochemically tracing nascent transcripts to infer synthesis and degradation rates in HEK293 cells upon CRISPRi of 180 genes. I found changes in synthesis and degradation rate to be predominantly negatively correlated with gene length (Figure 2D,E; Extended Data Tables 7 and 8; Extended Data Figure 9). Curiously, effects on length-correlated transcript synthesis appeared particularly pronounced after loss-of-function of genes involved in translation (Extended Data Figure 10). Overall, loss-of-function of these 180 genes resembled GLTD in aging by originating during the transcript synthesis^7^. It is not known if there is a length-correlated compensation of transcript stability in aging or if modulation of proteostasis^22^ affects GLTD in aging.

My findings align well with wear-and-tear-based theories of biological aging, especially imperfection-driven non-random damage^23^ and DNA damage^3,24^, but cannot exclude additional explanations and covariates or extend beyond the transcriptome. The main contribution of my present analysis is the demonstration that the primary^5^ transcriptional phenotype encountered during aging is a frequent mode of transcriptional change following perturbations. These transcriptional data suggest the hypothesis that “aging” is not a specific biological process but rather that during the passage of time, exposure to myriad perturbations accumulate in a manner that would be described as “aging”. Alternatively, the transcriptional phenotype of older individuals may reflect a higher susceptibility of their transcriptomes toward change following perturbations.

## Acknowledgments

I thank Zihan Xu and Junyue Cao for sharing intermediate data behind their publication. I thank Ben D. Singer and M. Jamal Jenkins for feedback on the manuscript.

## Author contributions

T.S. conceptualized the study, curated the data, performed formal analysis, visualized the findings, and wrote the manuscript.

## Materials and Methods

### Length Correlations

The primary transcript lengths for humans and mice were obtained from BioMart (http://useast.ensembl.org/biomart/martview), specifically from GRCh38.p13, and GRCm39, respectively. For each gene, the median length across all primary transcripts was calculated, and genes marked as protein-coding by NCBI’s reference file (https://ftp.ncbi.nlm.nih.gov/gene/DATA/GENE_INFO/All_Data.gene_info.gz) were selected. For mice and humans, only genes with an unambiguous 1:1 mapping between Entrez and Ensembl gene identifiers were retained. Length correlations were defined as the Spearman correlation between the lengths of primary transcripts and fold-changes of transcripts and computed using SciPy 1.2.1. P-values of Spearman correlations were obtain by Kendall and Stuart’s method for P-values of Spearman correlations ^11^.

### Bootstrapping

Bootstrapping was done with 100,000 iterations.

### Reanalysis of EBI-GXA

Expression profiles were downloaded from http://ftp.ebi.ac.uk/pub/databases/microarray/data/atlas/experiments/. To account for some experiments reporting few genes, experiments reporting less than 10,000 protein-coding genes with unambiguous 1:1 mapping between Entrez and Ensembl were excluded from the analysis.

### Reanalysis of GPSAdb

Expression profiles were downloaded from https://www.gpsadb.com using their bulk-download option. Length correlations were computed over the transcripts of genes that were not directly targeted by a given perturbation.

### Exclusion of pathways associated with aging

Gene Ontology annotations from NCBI were used, https://ftp.ncbi.nlm.nih.gov/gene/DATA/gene2go.gz. Genes were excluded if “go_term” matched case-insensitively the following regular expression, which represents the list of age-associated pathways compiled by Stegeman and Weake^17^: dna repair|inflamm|calcium-mediated signaling|\btranslation|insulin|response to heat|heat shock|cell cycle|protein catabolic process|\bliver|MAPK cascade|mitochondrial matrix|potassium|ATPase|glucose|apoptosis|retinoic|matrix|oxidative stress|RNA processing|cell differentiation|\bT cell activation|cellular matrix|metabolic process|axon guidance|synaptic trans|stress

### Reanalysis of Perturb-seq

The fold-changes and coordinates of Figure 2 of Replogle and Saunders et al.^20^ were obtained from the author-provided files at https://doi.org/10.25452/figshare.plus.21632564.v1.. Length correlations were computed over the transcripts of genes that were not directly targeted by a given perturbation. PCA analysis was preceded by an imputation of missing values according to the median fold-change observed for a gene, and z-scoring across of a gene over all perturbations.

### Transcription factor analysis

Transcription factor binding sites were obtained from GTRD^25^, version 21, http://gtrd.biouml.org:8888/downloads/current/intervals/target_genes/, and the included “genes promoter[-1000,+100]” used to map transcription factors toward genes.

### Gene Ontology enrichment analysis

Gene Ontology annotations from NCBI were used, https://ftp.ncbi.nlm.nih.gov/gene/DATA/gene2go.gz. Enrichment was done in Python via Fisher’s Exact test, implemented in scipy.stats.

### Reanalysis of Xu et al

Per-gene resolved intermediate data behind Xu et al.^21^ was requested from, and kindly shared by, the authors, Zihan Xu and Junyue Cao. Length correlations were computed over the transcripts of genes that were not directly targeted by a given perturbation.

## Data and code availability

Code will be made public upon acceptance via GitHub, and is meanwhile attached for reviewers as a zip file.

## Figures

**Extended Data Figure 1.**
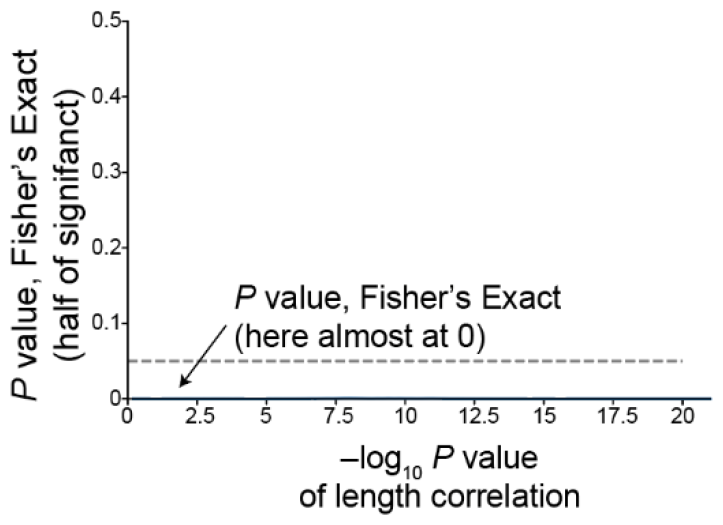
*P* value of two-sided Fisher’s Exact test for hypothesis that at a given significance threshold of length-correlated change (*P* value of length correlation) within EBI-GXA there was the same amount of negative and positive correlations.

**Extended Data Figure 2.**
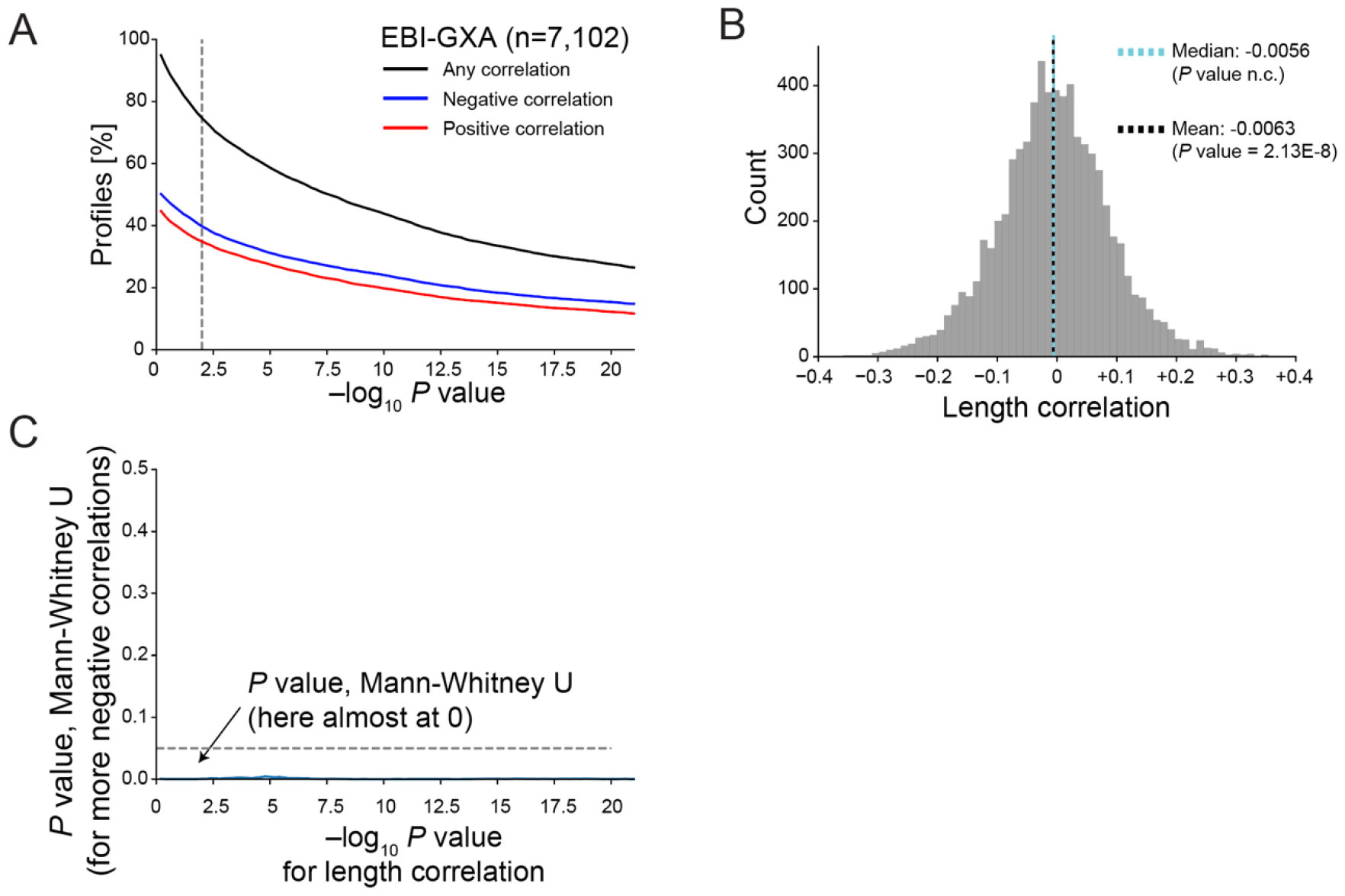
Gene Length-dependent Transcription Decline (GLTD) persists in EBI-GXA when excluding genes annotated for aging-related pathways. (A) Fraction of gene expression profiles with length-correlated change in EBI-GXA^10^ at different *P* value thresholds, determined by Kendall and Stuart’s method^11^. Dashed vertical line indicates *P* value of 0.01. (B) Histogram of all observed length-correlated changes in EBI-GXA, including non-significant ones. *P* value of median inferred from bootstrapping and *P* value of mean from two-sided one group T-test. n.c. is non computable as close to 0. (C) *P* value of two-sided Fisher’s Exact test for hypothesis that at a given significance threshold of length-correlated change (*P* value of length correlation) within EBI-GXA there was the same amount of negative and positive correlations.

**Extended Data Figure 3.**
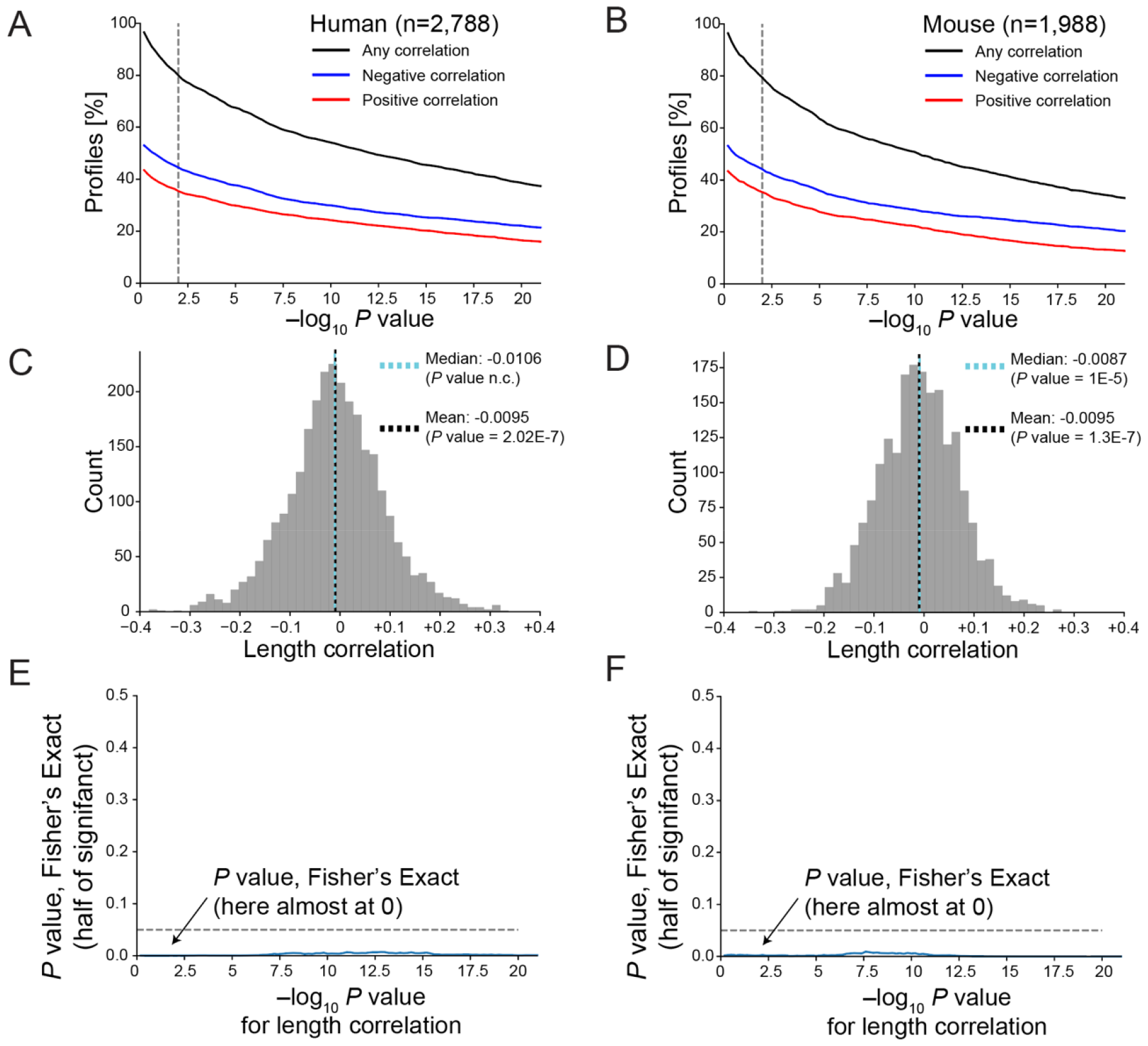
Gene Length-dependent Transcription Decline (GLTD) persists in EBI-GXA when only considering experiments annotated to have been performed by microarrays. (A) Fraction of gene expression profiles with length-correlated change for human studies at different *P* value thresholds, determined by Kendall and Stuart’s method^11^. Dashed vertical line indicates *P* value of 0.01. (B) As (A) but for mouse studies. (C) Histogram of all observed length-correlated changes in human studies, including non-significant ones. *P* value of median inferred from bootstrapping and *P* value of mean from two-sided one group T-test. n.c. is non computable as close to 0. (D) As (C) but for mouse studies. (E) *P* value of two-sided Fisher’s Exact test for hypothesis that at a given significance threshold of length-correlated change (*P* value of length correlation) within human studies there was the same amount of negative and positive correlations. (F) As (E) but for mouse studies.

**Extended Data Figure 4.**
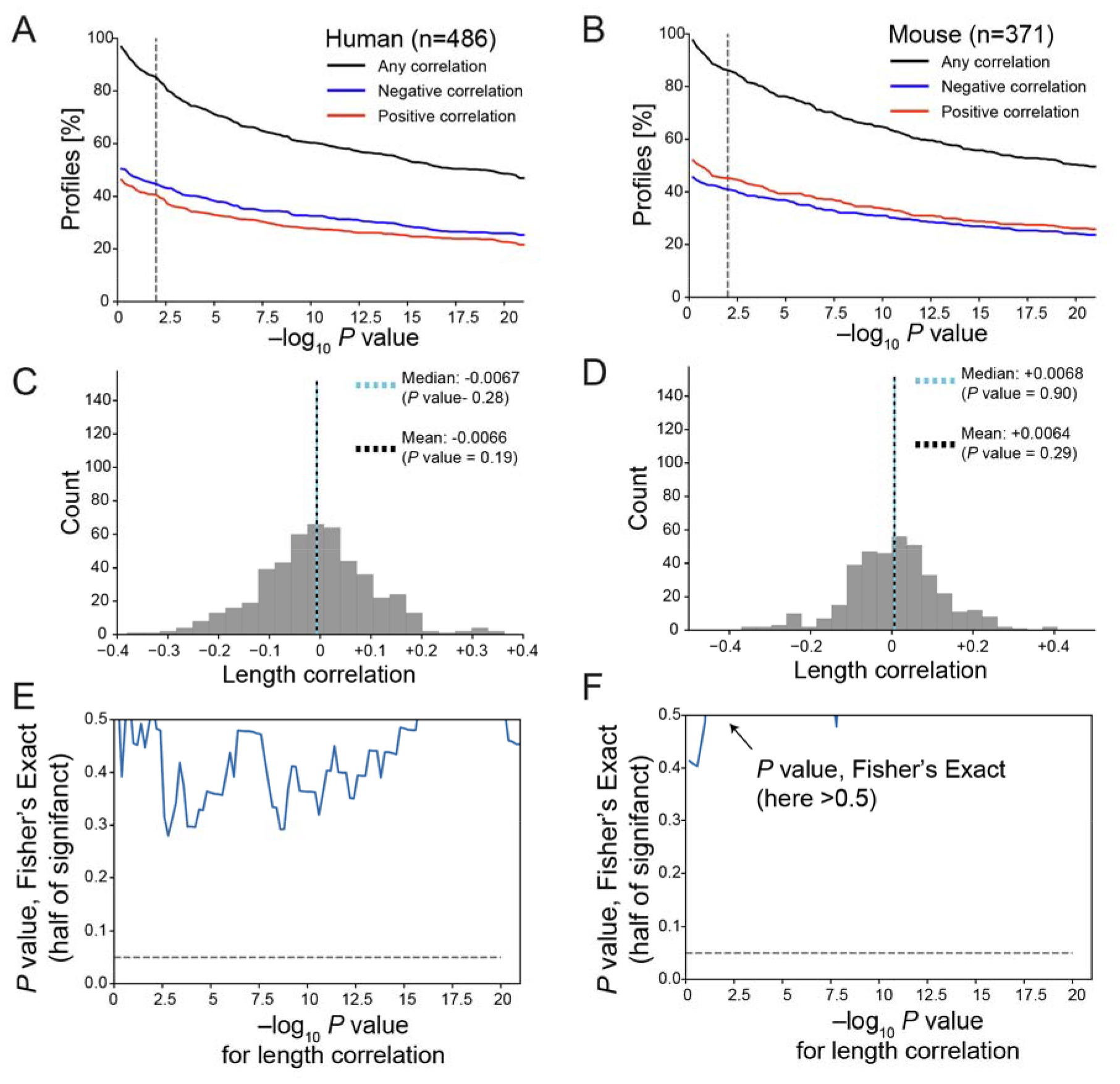
Gene Length-dependent Transcription Decline (GLTD) persists in EBI-GXA when only considering experiments annotated to have been performed by RNA-sequencing (RNA-Seq). (A) Fraction of gene expression profiles with length-correlated change for human studies at different *P* value thresholds, determined by Kendall and Stuart’s method^11^. Dashed vertical line indicates *P* value of 0.01. (B) As (A) but for mouse studies. (C) Histogram of all observed length-correlated changes in human studies, including non-significant ones. *P* value of median inferred from bootstrapping and *P* value of mean from two-sided one group T-test. n.c. is non computable as close to 0. (D) As (C) but for mouse studies. (E) *P* value of two-sided Fisher’s Exact test for hypothesis that at a given significance threshold of length-correlated change (*P* value of length correlation) within human studies there was the same amount of negative and positive correlations. (F) As (E) but for mouse studies.

**Extended Data Figure 5.**
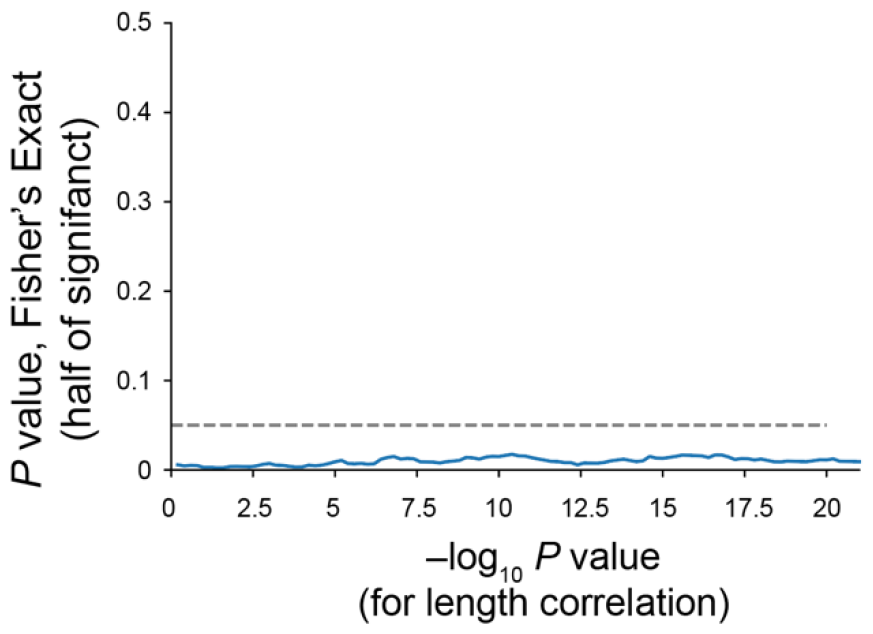
*P* value of two-sided Fisher’s Exact test for hypothesis that at a given significance threshold of length-correlated change (*P* value of length correlation) within GPSAdb there was the same amount of negative and positive correlations.

**Extended Data Figure 6.**
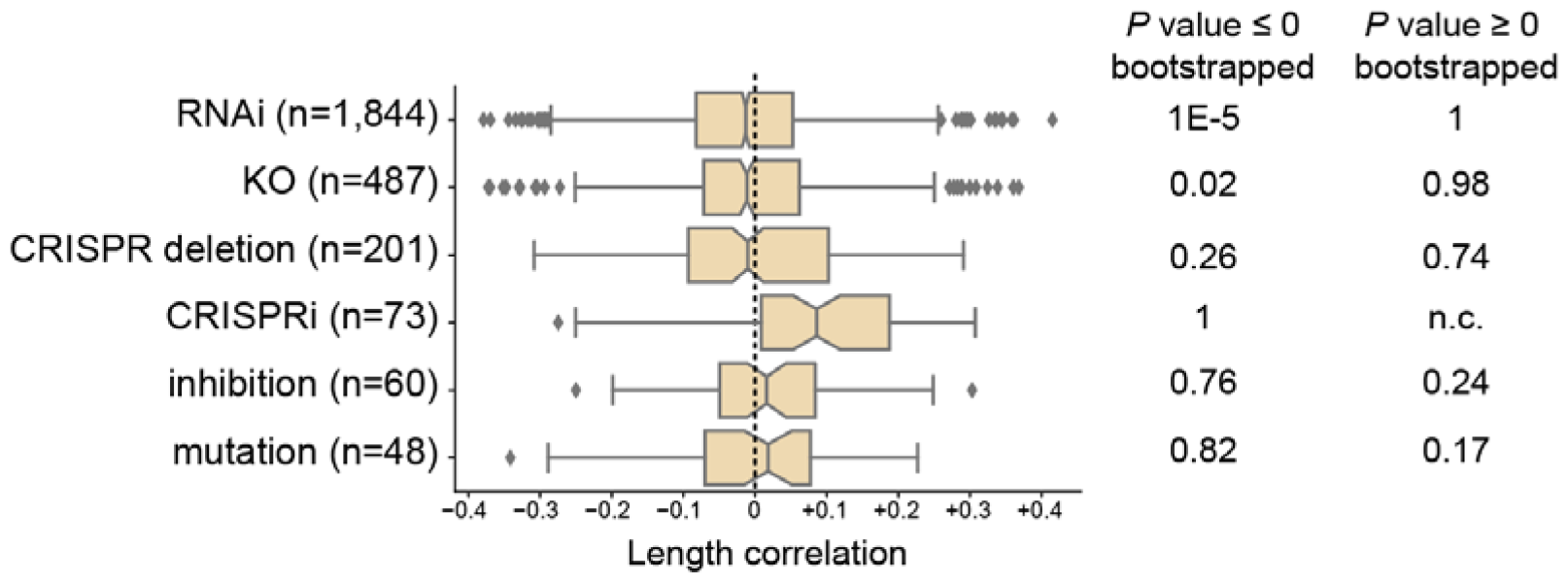
Length correlations within GPSAdb according to method used for loss-of-function. Notches indicate bootstrapped 95% confidence interval of median. Box contains 25-75% range of observe length correlations. n indicates number of genes. n.c. indicates that not precisely computable.

**Extended Data Figure 7.**
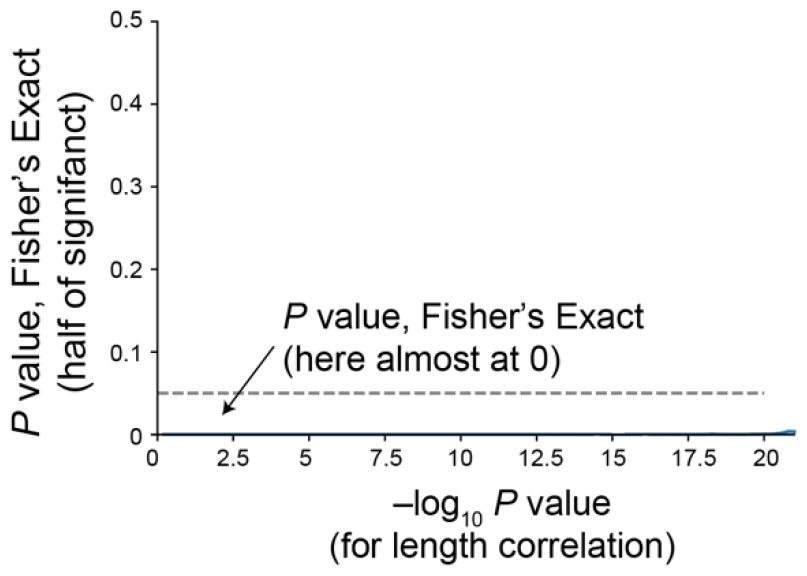
*P* value of two-sided Fisher’s Exact test for hypothesis that at a given significance threshold of length-correlated change (*P* value of length correlation) within Replogle et al.^20^ there was the same amount of negative and positive correlations.

**Extended Data Figure 8.**
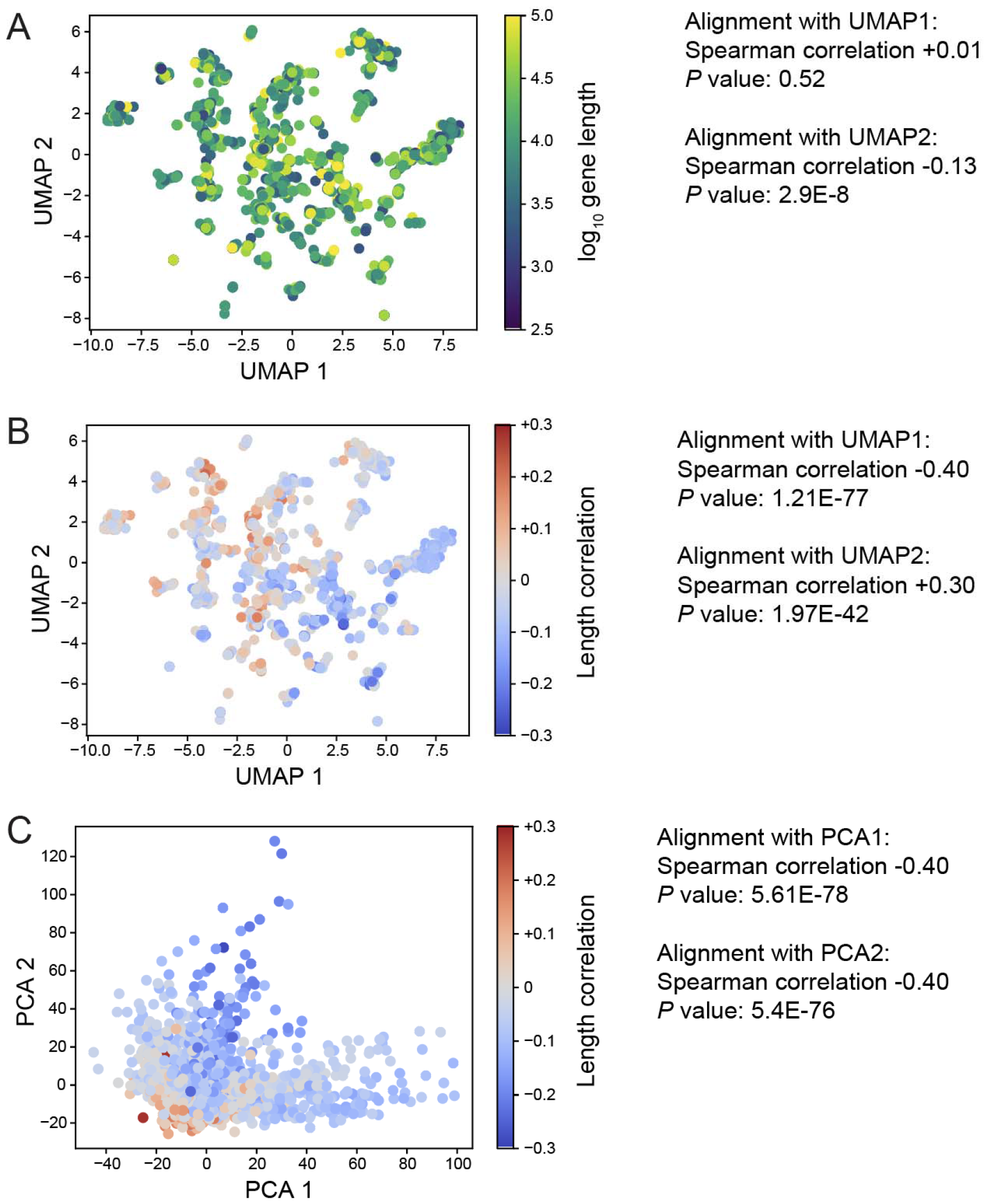
The alignment of length-correlated change with the map of loss-of-function effects by Replogle et al.^20^ is robust toward alternative analysis. (A) Superimposition of gene-length of targeted genes (dots). (B) Superimposition of length-correlated change computed without genes annotated for pathways associated with aging. (C) Alternative map of loss-of-function using Principal Component Analysis (PCA).

**Extended Data Figure 9.**
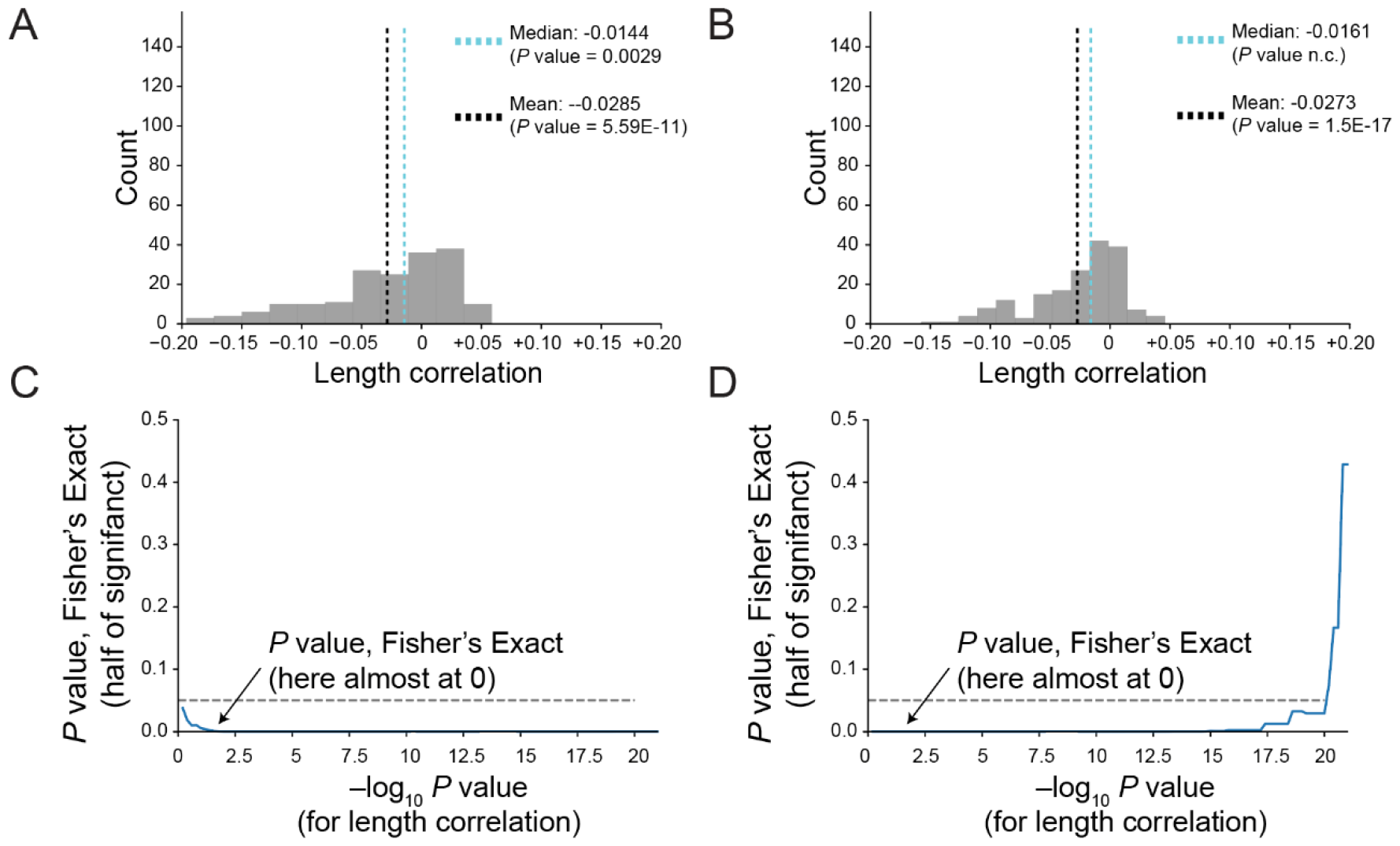
Loss-of-function causes length-correlated change of transcript synthesis and degradation when re-analyzing Xu et al.^21^. (A) Histogram of all observed length-correlated changes in transcript synthesis Xu et al., including non-significant ones. *P* value of median inferred from bootstrapping and *P* value of mean from two-sided one group T-test. n.c. is non computable as close to 0. (B) As (A) but for degradation rates. (C) *P* value of two-sided Fisher’s Exact test for hypothesis that at a given significance threshold of length-correlated change (*P* value of length correlation) within transcriptome synthesis data of Xu et al. there was the same amount of negative and positive correlations. (D) Same as (C) but for degradation rates.

**Extended Data Figure 10.**
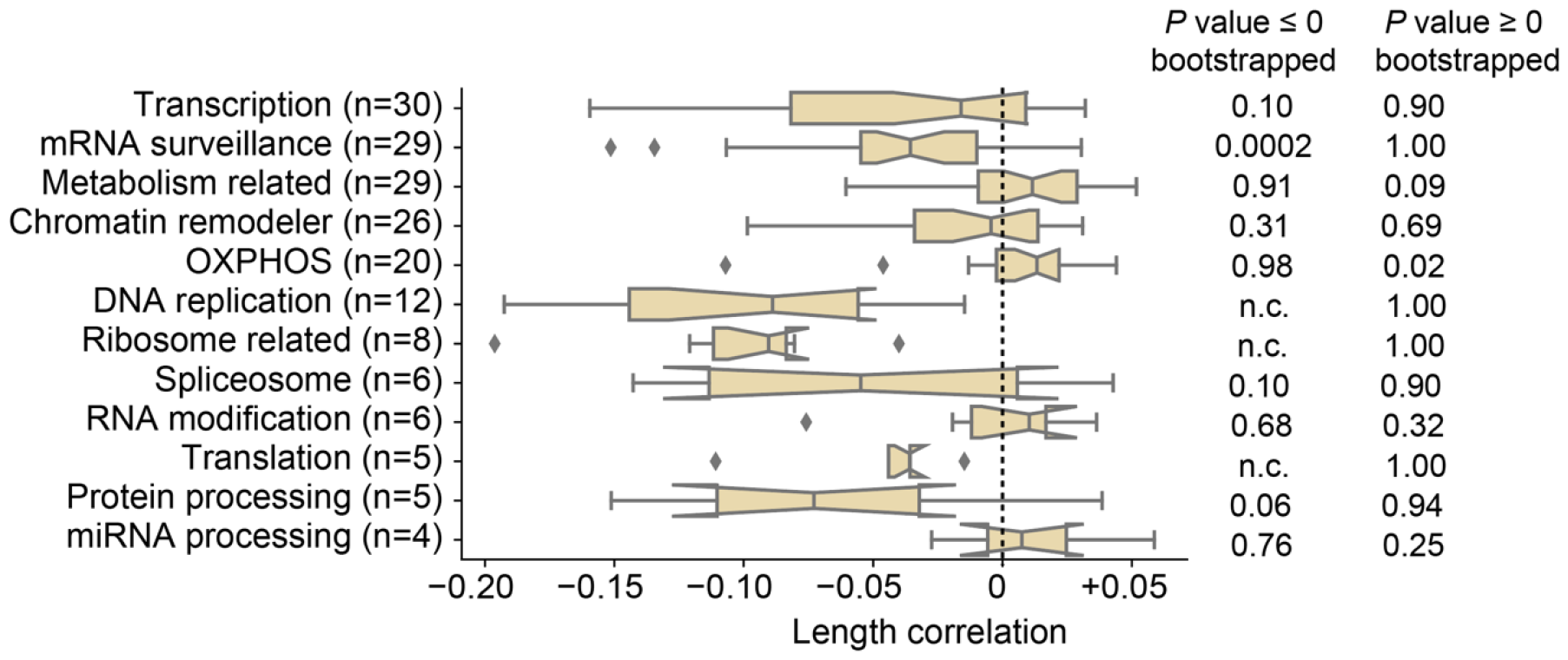
Length correlations within Xu et al.^21^ according to process annotations by them. Notches indicate bootstrapped 95% confidence interval of median. Box contains 25-75% range of observe length correlations. n indicates number of genes.

**Extended Data Table 1**. Manual inspection of 10 randomly selected gene expression profiles in EBI-GXA.

**Extended Data Table 2**. Length-correlated changes in EBI-GXA. Rho is Spearman correlation, pval the *P* value for correlation obtained by Kendall and Stuart’s method^11^. n is number of protein-coding genes.

**Extended Data Table 3**. Length-correlated changes in GPSAdb. Rho is Spearman correlation, pval the *P* value for correlation obtained by Kendall and Stuart’s method^11^. n is number of protein-coding genes.

**Extended Data Table 4**. Length-correlated changes in Replogle et al.^20^. Rho is Spearman correlation, pval the *P* value for correlation obtained by Kendall and Stuart’s method^11^. umap_x is first UMAP dimension, and umap_y is the second UMAP dimension.

**Extended Data Table 5**. Associations of transcription factors with first and second UMAP dimension (axis). Correlations obtained via point-biserial correlations and Spearman correlations.

**Extended Data Table 6**. Gene Ontology analysis for genes with GLTD in Replogle et al.^20^.

**Extended Data Table 7**. Length-correlated changes for synthesis in Xu et al.^21^. Rho is Spearman correlation, pval the *P* value for correlation obtained by Kendall and Stuart’s method^11^. gene_class is annotation of process provided by authors.

**Extended Data Table 8**. Length-correlated changes for degradation in Xu et al.^21^. Rho is Spearman correlation, pval the *P* value for correlation obtained by Kendall and Stuart’s method^11^. gene_class is annotation of process provided by authors.

## References

1 Kennedy, B. K. et al. Geroscience: linking aging to chronic disease. Cell 159, 709–713 (2014). 10.1016/j.cell.2014.10.039

2 Freund, A. Untangling Aging Using Dynamic, Organism-Level Phenotypic Networks. Cell Syst 8, 172–181 (2019). 10.1016/j.cels.2019.02.005

3 Vermeij, W. P. et al. Restricted diet delays accelerated ageing and genomic stress in DNA-repair-deficient mice. Nature 537, 427–431 (2016). 10.1038/nature19329

4 Hall, H. et al. Transcriptome profiling of aging Drosophila photoreceptors reveals gene expression trends that correlate with visual senescence. BMC Genomics 18, 894 (2017). 10.1186/s12864-017-4304-3

5 Stoeger, T. et al. Aging is associated with a systemic length-associated transcriptome imbalance. Nature Aging 2, 1191–1206 (2022). 10.1038/s43587-022-00317-6

6 Ibañez-Solé, O., Barrio, I. & Izeta, A. Age or lifestyle-induced accumulation of genotoxicity is associated with a length-dependent decrease in gene expression. iScience, 106368 (2023). 10.1016/j.isci.2023.106368

7 Gyenis, A. et al. Genome-wide RNA polymerase stalling shapes the transcriptome during aging. Nat Genet 55, 268–279 (2023). 10.1038/s41588-022-01279-6

8 Soheili-Nezhad, S., Ibáñez-Solé, O., Izeta, A., Hoeijmakers, J. H. & Stoeger, T. Time is ticking faster for long genes in aging. Trends in Genetics (in press).

9 Hartmann, A. et al. Ranking Biomarkers of Aging by Citation Profiling and Effort Scoring. Front Genet 12, 686320 (2021). 10.3389/fgene.2021.686320

10 Papatheodorou, I. et al. Expression Atlas: gene and protein expression across multiple studies and organisms. Nucleic Acids Res 46, D246–D251 (2018). 10.1093/nar/gkx1158

11 Kendall, M. G. & Stuart, A. Inference and Relationship. The Advanced Theory of Statistics 2 (1973).

12 Niedernhofer, L. J. et al. A new progeroid syndrome reveals that genotoxic stress suppresses the somatotroph axis. Nature 444, 1038–1043 (2006). 10.1038/nature05456

13 Cellerino, A. & Ori, A. What have we learned on aging from omics studies? Semin Cell Dev Biol 70, 177–189 (2017). 10.1016/j.semcdb.2017.06.012

14 Schaum, N. et al. Ageing hallmarks exhibit organ-specific temporal signatures. Nature 583, 596–602 (2020). 10.1038/s41586-020-2499-y

15 Poganik, J. R. et al. Biological age is increased by stress and restored upon recovery. Cell Metabolism 35, 807-+ (2023). 10.1016/j.cmet.2023.03.015

16 Gene Ontology, C. Gene Ontology Consortium: going forward. Nucleic Acids Res 43, D1049–1056 (2015). 10.1093/nar/gku1179

17 Stegeman, R. & Weake, V. M. Transcriptional Signatures of Aging. J Mol Biol 429, 2427–2437 (2017). 10.1016/j.jmb.2017.06.019

18 Mandelboum, S., Manber, Z., Elroy-Stein, O. & Elkon, R. Recurrent functional misinterpretation of RNA-seq data caused by sample-specific gene length bias. PLoS Biol 17, e3000481 (2019). 10.1371/journal.pbio.3000481

19 Guo, S. et al. GPSAdb: a comprehensive web resource for interactive exploration of genetic perturbation RNA-seq datasets. Nucleic Acids Res 51, D964–D968 (2023). 10.1093/nar/gkac1066

20 Replogle, J. M. et al. Mapping information-rich genotype-phenotype landscapes with genome-scale Perturb-seq. Cell 185, 2559–2575 e2528 (2022). 10.1016/j.cell.2022.05.013

21 Xu, Z., Sziraki, A., Lee, J., Zhou, W. & Cao, J. Dissecting key regulators of transcriptome kinetics through scalable single-cell RNA profiling of pooled CRISPR screens. Nature Biotechnology (2023). 10.1038/s41587-023-01948-9

22 Labbadia, J. & Morimoto, R. I. The biology of proteostasis in aging and disease. Annu Rev Biochem 84, 435–464 (2015). 10.1146/annurev-biochem-060614-033955

23 Gladyshev, V. N. The origin of aging: imperfectness-driven non-random damage defines the aging process and control of lifespan. Trends Genet 29, 506–512 (2013). 10.1016/j.tig.2013.05.004

24 Schumacher, B., Pothof, J., Vijg, J. & Hoeijmakers, J. H. J. The central role of DNA damage in the ageing process. Nature 592, 695–703 (2021). 10.1038/s41586-021-03307-7

25 Yevshin, I., Sharipov, R., Kolmykov, S., Kondrakhin, Y. & Kolpakov, F. GTRD: a database on gene transcription regulation-2019 update. Nucleic Acids Res. 47, D100–D105 (2019). 10.1093/nar/gky1128

